# *Mysm1* mutations in *meander tail* mice cause anterior-selective cerebellum malformation

**DOI:** 10.64898/2026.02.25.708017

**Authors:** Bruce A. Hamilton, Dorothy Concepcion, Max Chang, Christopher Benner, Catherine Liang, Nathan R. Zemke, Melissa Gymrek, Dan Goldowitz, Colin Fletcher

## Abstract

Mouse *meander tail* (*mea*) mutations produce kinked tails and selective malformation of the cerebellum anterior compartment. The anterior cerebellum defects are cell autonomous with respect to granule cell precursors, but the molecular basis has not been known. Myb-like, SWIRM, and MPN domain containing protein 1 (MYSM1) is a chromatin-associated deubiquitinase that promotes gene expression by removing monoubiquitin from histone H2A, among other targets. Loss of MYSM1 function in mice or humans results in bone marrow failure with defective maturation of B cell lineages. Here we show that extant *mea* alleles have mutations in *Mysm1* and cause both neurological and hematological phenotypes, as do new non-complementing endonuclease-mediated mutations. Multimodal single-nucleus assays show *Mysm1* effects on gene expression in several lineages and on the proportion of granule cell precursors by E14.5. Intriguingly, *Mysm1* orthologs have been independently lost in several animal and fungal lineages, including yeast, flies, and nematodes. These results unite previously disconnected literature and demonstrate a requirement for MYSM1 activity in compartment-specific development of the cerebellum and suggest potential for compensatory pathways.

**Significance Statement:** Perturbations to core regulatory machinery often produce pleiotropic effects and even intensively studied systems can have significant phenotypic effects that were not assessed in models developed for a different purpose. Here we show that classical *meander tail* mice, characterized by ankylosing spondylitis in tail vertebrae and a cerebellum malformation that defined the anterior-posterior compartment boundary, have mutations in *Mysm1*, encoding a histone 2A deubiquitinase. We show pigmentation defects and hematopoietic abnormalities that model human disease. While *Mysm1* mutations change gene expression patterns in many cerebellar cell types, they selectively decrease the proportion of granule cell lineages. Recurrent loss of *Mysm1* orthologs across fungal and animal phylogenies suggests the potential for bypass mechanisms.

## Introduction

*Meander tail* (*mea*) was first described nearly 50 years ago as a spontaneous mutation with variably kinked tail and unsteady gait [1]. Subsequent studies of *mea* mice identified a cerebellar malformation that defined a discrete anterior-posterior developmental compartment boundary, with selective hypoplasia of the anterior lobes [2, 3] resulting from a cell-autonomous defect in granule cell precursors [4–6]. Despite substantial interest, *mea* alleles were never identified with a molecular cause. Only the eponymous kinked tail in published descriptions of *mea* overlap reported phenotypes for *Mysm1* knockout mice, which showed more severe tail phenotypes than those noted for *mea*.

*Mysm1* encodes a conserved deubiquitinase associated with both chromatin and cytoplasmic targets. MYSM1 is named for its protein domain architecture, comprising a <u>My</u>b-like (SANT) domain, a <u>S</u>WIRM domain, and a <u>M</u>PN+ proteolytic domain that includes a zinc-coordinating JAMM motif required for isopeptidase activity. MYSM1 was first identified as a deubiquitinase for histone H2A (2A-DUB) monoubiquitinated at lysine 119 (H2AK119Ub) that regulated transcription initiation and elongation by coordinating H2A ubiquitination and acetylation and destabilizing the H1 linker histone, in complexes containing EP300 and PCAF (KAT2B) [7, 8]. Knockout mouse experiments showed that *Mysm1* was essential for hematopoiesis and *Ebf1* activation in early B-cell lineage, while also showing truncated tails, abdominal white spotting due to defects in melanoblast migration, and tremor and ataxia with unclear cellular bases [9–11]. In addition to nuclear roles, cytoplasmic MYSM1 down-regulates innate immunity through inactivation of TRAF3 and TRAF6 [12]. Following the initial mouse models, homozygous *MYSM1* mutations were identified in consanguineous human subjects with severe anemia, bone marrow failure that included high genotoxic stress in hematopoietic progenitors, neurodevelopmental delay, and skeletal abnormalities [13–15]. Cerebellar abnormalities related to MYSM1 activity have not been previously reported.

Here we show that the extant classical *mea* alleles (*mea^J^*and *mea^2J^*), which arose close in time in the same Jackson Laboratory production colony (BKS.Cg– + *Lepr^db^* / *Dock7^m^*+) [4, 16, 17], are distinct alleles of *Mysm1*. Both *mea^J^* and *mea^2J^* produced abdominal white spotting and reduced blood cell counts consistent with *Mysm1* but not previously reported for *mea* mutations. New endonuclease-mediated *Mysm1* alleles failed to complement *mea*, confirming these *Mysm1* variants as the causal mutations for both the vertebral malformation and the anterior cerebellum defect. Combined single-nucleus RNA- and ATAC-seq from E14.5 cerebellum for two mutant alleles compared with littermates showed genotype-dependent gene expression changes across multiple cell types. While granule cell precursors (GCPs) had comparatively fewer differentially expressed genes, the proportion of GCPs was selectively reduced in mutants across four littermate pairs. Surprisingly, while phylogenetic analysis found MYSM1 orthologs in animals, fungi, and Amoebozoa, the gene has been recurrently lost from lineages represented by common laboratory organisms–including yeasts, flies, and nematodes. In resolving a nearly 50-year-old mystery for the basis of *mea* mutations, these results unite previously separate literatures on developmental neurobiology in *mea* mice and gene regulatory activity by MYSM1 in both health and disease.

## Results

### Distinct *Mysm1* variants in *mea* alleles

Linkage mapping localized the *mea* locus within chromosome 4, between *Ifa* and *Glut1*, in a backcross of *mea^2J^* with CAST/EiJ partners [18]. A previously unpublished extension of this cross implicated an interval proximal to the 5’ end of *Nfia* but with an uncertain proximal boundary, likely due to incomplete penetrance in this intersubspecific cross. To narrow the interval, we added intercross progeny from two standard laboratory strains and *mea^J^*, reasoning that partner strains with different *Prdm1* alleles and an alternate *mea* allele might increase the diversity of recombination breakpoints [19, 20], improve penetrance, and/or allow progeny testing of any conflicting recombinants. Combined data placed *mea* between 91.0 and 95.7 Mb of the mm39 mouse reference genome (Figure 1A).

**Figure 1.**
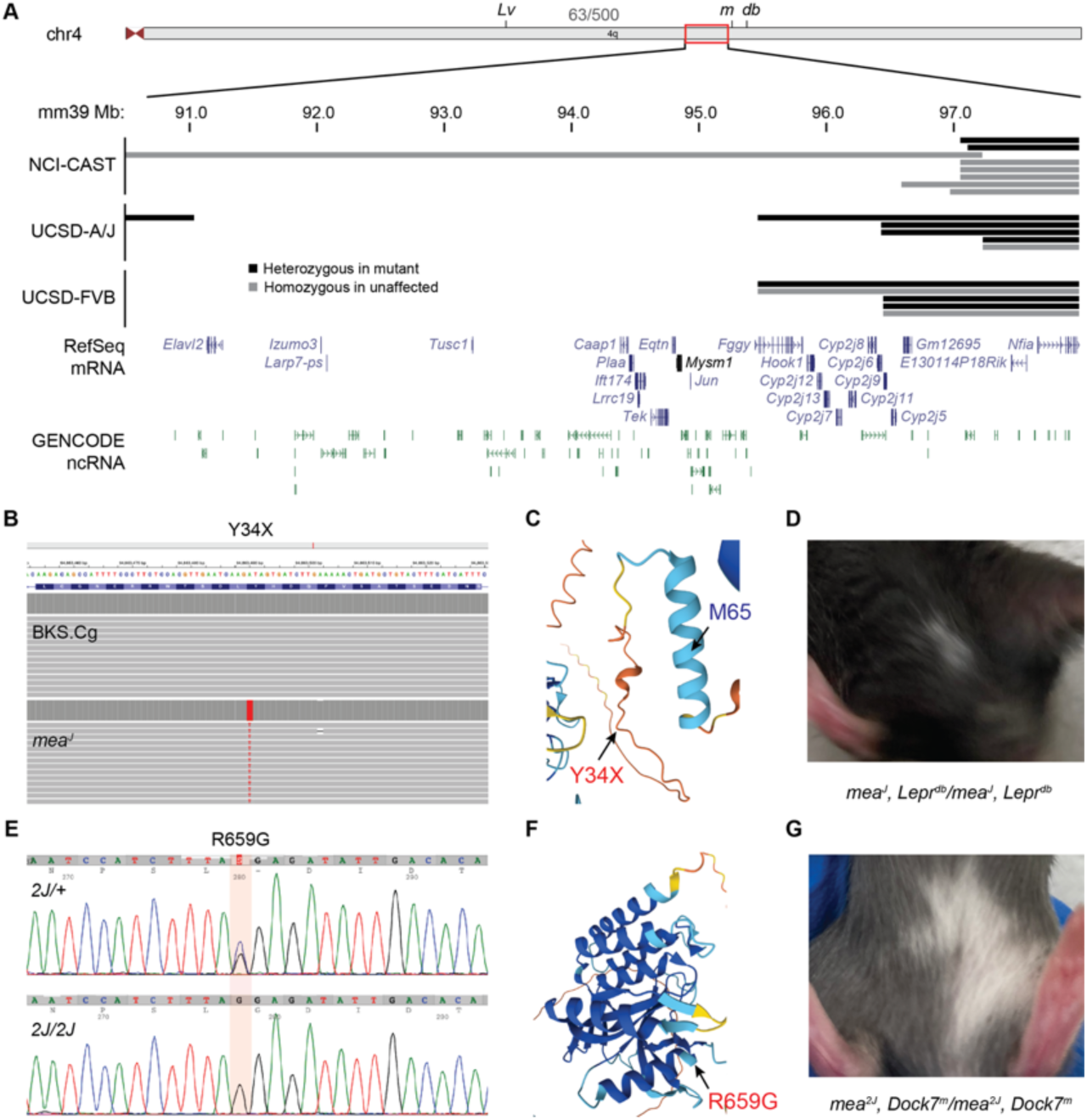
Distinct *Mysm1* mutation in each *mea* allele. (**A**) Genetic crosses localized *mea* alleles on chromosome 4. Regions definitively excluded by heterozygosity in an affected animal shown as black bars. Recombinants whose interpretation relied on penetrance in an unaffected animal are shown in grey. (**B**) PacBio HiFi reads identified a Y34X variant in *mea^J^* not present in its BKS.Cg parental strain control. (**C**) AlphaFold2 model of mouse MYSM1 shows position of the Y34X nonsense variant and the next in-frame methionine at residue 65. (**D**) Small patches of abdominal white spotting were seen on most BKS.Cg–*mea^J^* homozygotes. (E) Sanger sequencing of *Mysm1* exons identified a R659G missense variant in *mea^2J^*. (F) AlphaFold2 model shows position of the R659G variant on the surface of the catalytic MPN domain. (G) Larger patches of white spotting were typical of BKS.Cg–*mea^2J^* homozygotes.

DNA sequencing identified distinct *Mysm1* mutations in *mea^J^* and *mea^2J^* within this interval. We first performed whole genome, high fidelity, long-read sequencing to identify variants in the interval and assess the background polymorphism rate between *mea^J^* mutant and its BKS.Cg–*Dock7^m^ + / + Lepr^db^* parental strain (PacBio Revio platform, 15X coverage for each strain). Complete sequence coverage for the *mea* interval identified a single sequence variant within the interval that distinguished *mea^J^*from both its BKS.Cg parental stock and the C57BL/6J reference genome: a T>A transversion on the *Mysm1* coding strand, creating a Y34X nonsense mutation in exon 2 relative to reference sequence NP_796213.2 (Figure 1B). The occurrence of a potential reinitiation codon at position M65 (Figure 1C) raised the possibility that this variant may not be null [21–23]. Consistent with *Mysm1* knockout mice, but not previously noted for *mea*, we observed small white spots on the abdominal fur of most *mea^J^* homozygotes (Figure 1D). The *mea^2J^* allele was identified in the same BKS.Cg stock [16, 17]. PCR-based sequencing of all 20 exons identified a single variant, a C>G transversion in exon 16 (Figure 1E), encoding a R659G amino acid substitution at a highly conserved residue within the JAMM isopeptidase motif of the MPN domain (Figure 1F), for which homologous position encoded arginine in all 467 mammals with data in the MultiZ alignment track of the UCSC Genome Browser [24]. Abdominal spotting was typically more prominent in *mea^2J^* than *mea^J^* (Figure 1G). Equivalent variants in human *MYSM1* (p.Y37X and p.R668G relative to NP_001078956) are each observed once in the gnomAD 4.1 database [25]. Data in gnomAD argue against *MYSM1* being heterozygous loss-of-function intolerant in humans (pLI 0.09, LOEUF 0.578), consistent with biallelic inactivation in bone marrow failure syndrome [13, 15]. Both equivalent human variants are predicted deleterious by multiple variant effect prediction tools [26].

### Edited mutations in *Mysm1* are allelic to *meander tail*

We confirmed that *Mysm1* variants in *mea^J^* and *mea^2J^* are causal by complementation tests with new *Mysm1* alleles created by CRISPR/Cas9 editing in FVB/NJ mice. We recovered both a frame-shifting 23-bp deletion within exon 2, encoding a predicted D32Efs*13 variant relative to NP_796213.2, and a 96-bp deletion that spanned the splice donor site into intron 3 (Figure 2A). D32Efs*13 created an early premature termination codon similar to the Y34X mutation in *mea^J^* but with a shorter distance to a potential reinitiation at codon M65. Compound heterozygotes for *mea^J^* and Δ96 had shorter, kinked tails relative to either heterozygote, confirming allelism with respect to ankylosing spondylitis in tail vertebrae (Figure 2B). Both *mea* alleles and D32Efs*13 compound heterozygotes also recapitulated blood cell phenotypes reported in *Mysm1* knockout mice and *MYSM1* patients [10, 13], with reductions in white blood cells, neutrophils, lymphocytes, and red blood cells but increased in mean corpuscular volume and mean corpuscular hemoglobin (Figure 2C). *Mysm1* D32Efs*13 and *mea* variants were also allelic with respect to the anterior-selective cerebellum malformation (folia I-VIa, Figure 2D). Both D13Efs*13 homozygotes and compound heterozygotes had less pronounced reductions in anterior cerebellum than the spontaneous *mea* mutations (Figure 2E); this could reflect either allelic differences or the effect of a more robust FVB genetic background on size of cerebellar folia [27]. We also note that all alleles and compound heterozygotes included a more modest but reproducible reduction of the posterior compartment (Figure 2E). D32Efs*13 homozygotes and compound heterozygotes with *mea^J^* had normal tails, suggesting a hypomorphic presentation with respect to the ankylosing spondylitis underlying the *meander tail* vertebral phenotype.

**Figure 2.**
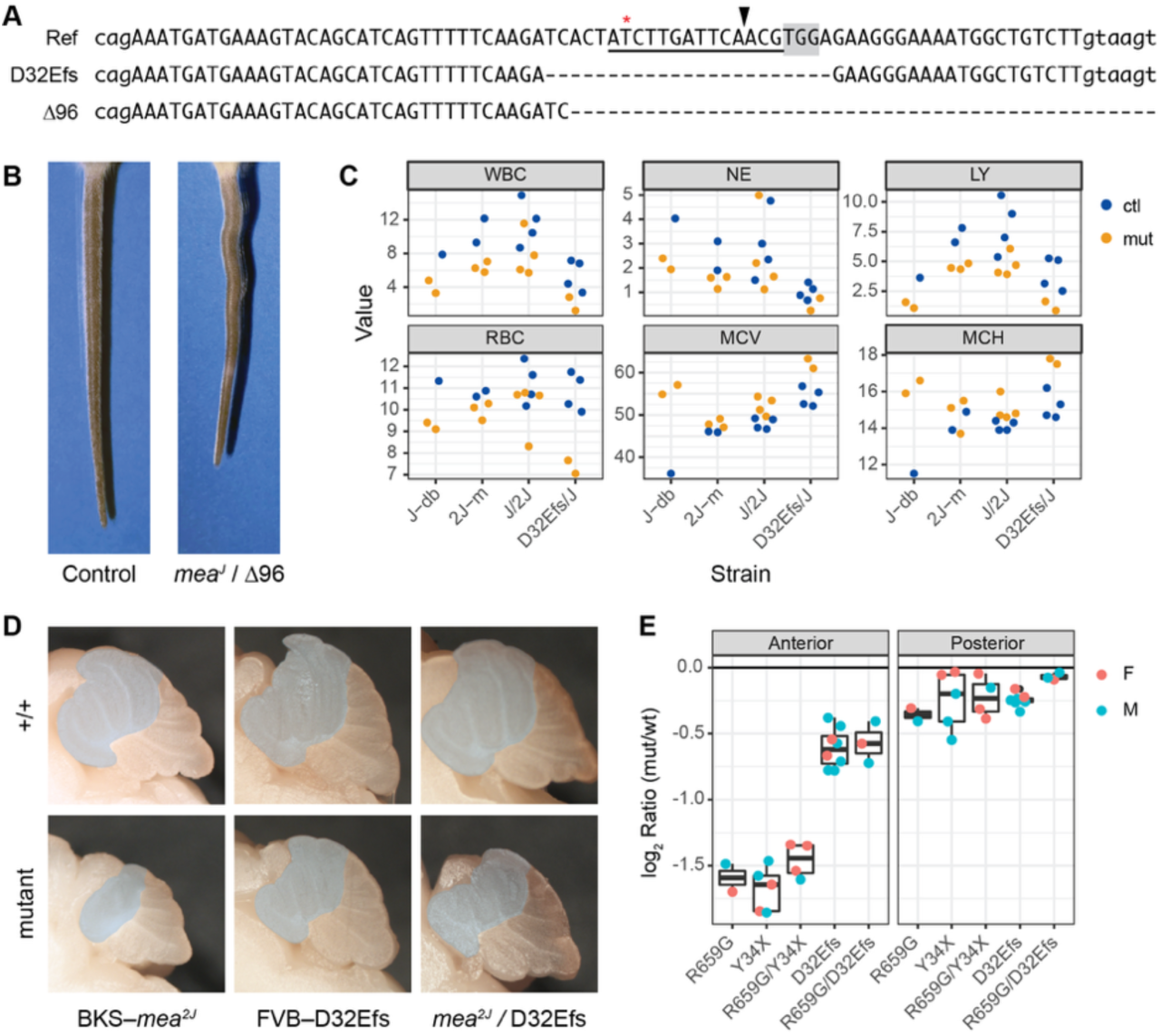
*Mysm1* edited variants are allelic to *mea*. (**A**) Reference (Ref) and 23-bp deletion allele (D32Efs) recovered at exon 2. RNA guide sequence is underlined, with protospacer-adjacent motif shaded. Arrowhead, predicted cleavage site. Lowercase intron, uppercase exon sequences. Asterisk, T>A in *mea^J^*. (**B**) Compound *mea^J^*/Δ96 heterozygotes had shorter, kinked tails with variegated pigmentation characteristic of *mea*. (**C**) Classical *mea* alleles (from stocks carrying closely-linked *Lepr^db^* or *Dock7^m^* mutation) and compound heterozygotes between *mea^J^* and either *mea^2J^* or D32Efs all showed reductions in white blood cells (WBC), neutrophils (NE), lymphocytes (LY), and red blood cells (RBC), but elevated mean corpuscular volume (MCV) and mean corpuscular hemoglobin (MCH), consistent with the bone marrow failure reported for *Mysm1* knockout mice. Mutants in orange, sex-matched littermate controls navy. (**D**) Midline sagittal images illustrate selective reduction of the anterior compartment (folia I-VIa, grey mask) in *mea^2J^*, D32Efs, and compound heterozygous mutations relative to littermate controls. (**E**) Measured areas at midline showed dramatic reductions of the anterior and slight reductions in the posterior cerebellum relative to sex-matched littermates in both females (orange) and males (cyan).

### *Mysm1* early premature termination codon variants are hypomorphic

Both Y34X (*mea^J^*) and D32Efs*13 (edited) mutations predict premature termination codons upstream of a potential reinitiation codon, M65 in exon 3 (Figure 3A), which would produce a 755 amino acid, ∼86.4 kD protein lacking the unstructured aminoterminal end and part of a predicted alpha helix (Figure 1C). Similar human variants have been reported in ClinVar (Figure 3B). Western blots from embryonic day 17.5 (E17.5) mouse brains showed reduced or absent MYSM1 protein levels in D32Efs*13, Y34X (*mea^J^*), and R659G (*mea^2J^*) relative to control littermates (Figure 3C) and a TUBG loading control (Figure 3D). Residual MYSM1 in D32Efs*13 showed a mobility shift equivalent to ∼8 kD. Residual MYSM1 protein expression explains the milder phenotypes from this allele (Figure 2). A much weaker band at this position was seen across genotypes, suggesting a potentially similar endogenous protein variant. Major MYSM1 reactivity was also lost in R659G homozygotes, suggesting that this substitution may be destabilizing in vivo. Single-vector dual luciferase reporter assays [21] further supported non-null mechanisms with a quantitative basis for both D32Efs*13 and Y34X early premature termination codon variants (Figure 3E), with reporter expression for each allele significantly reduced by mutation of potential reinitiation codon 65 from AUG to CUG (M65L; p = 0.03 for D32Efs*13, p = 0.0002 for Y34X, Tukey’s HSD test after ANOVA for all mouse variants tested). This is similar to our recent findings with *Zfp423* early premature termination codon mutations, where expression below Western blot detection threshold nonetheless retained modest translational potential and slightly reduced phenotypic severity [28].

**Figure 3.**
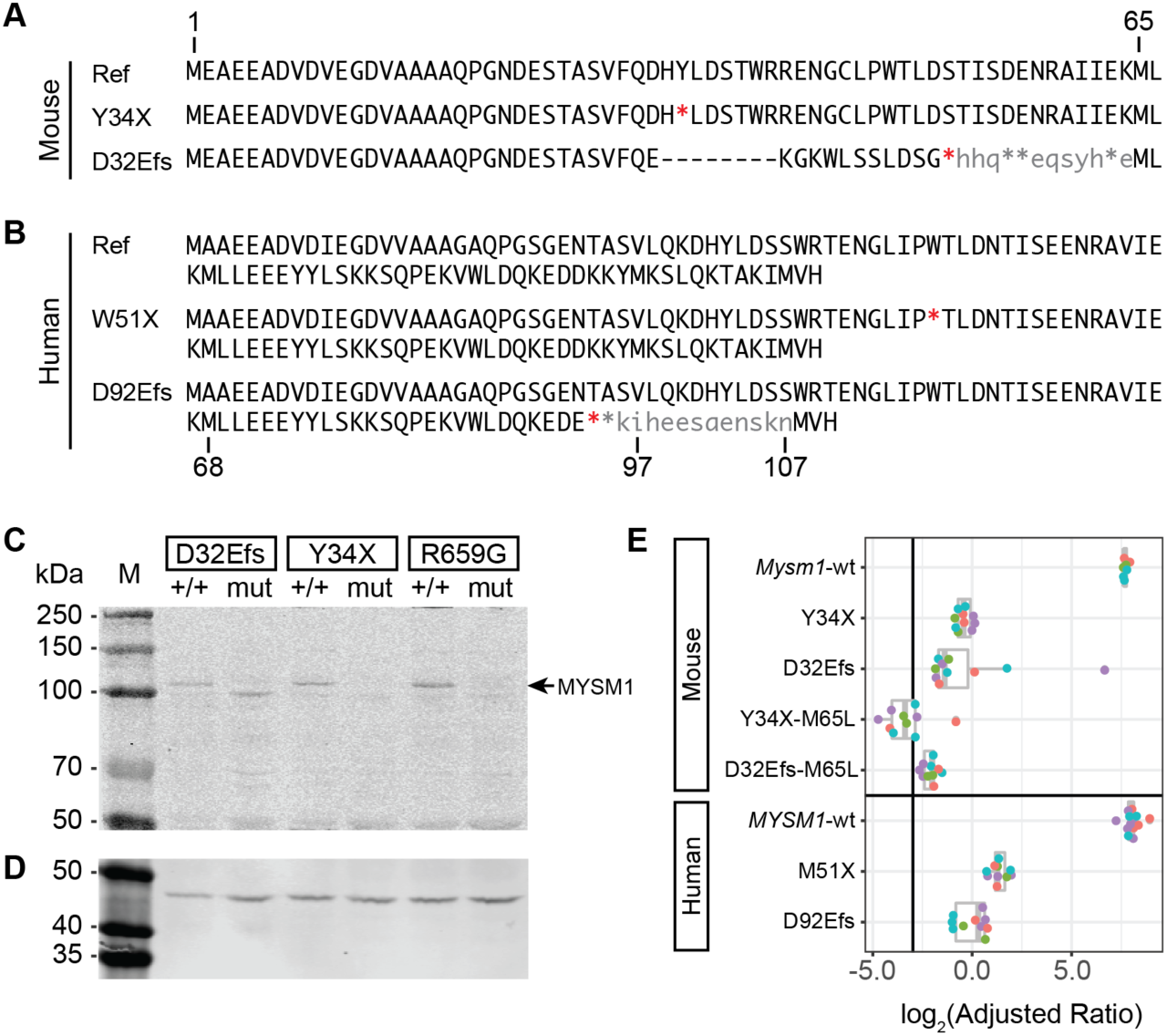
Y34X and D32Efs*13 allow residual MYSM1 protein expression. (**A**) Amino acid sequence for mouse reference, Y34X (*mea^J^*), and D32Efs*13 (Δ23) alleles, showing deleted (-), termination (*), and next available in-frame AUG (M65) codons. (**B**) Comparable human reference and ClinVar alleles W51X and D92Efs relative to potential reinitiation codons at M68, M97, and M107. (**C**) Western blots of E17.5 mouse brains from control (+/+) and mutant (mut) littermate pairs for each of the three *Mysm1/mea* mutations using rabbit anti-MYSM1 antibody. Arrow, wild-type protein size. (**D**) Loading control on the same membrane using mouse anti-TUBG. (**E**) Dual luciferase assays using aminoterminal fusions of the fragments shown in (A) for both mouse and human MYSM1 variants plotted as log2 of the ratio of Nano luciferase to firefly luciferase. Colors denote two to three replicate transfections for each of four independent plasmid preparations. Vertical line, median of three negative controls with a frameshifted Nano luciferase.

The divergence between D32Efs*13 and Y34X results in Western blots but not in the intronless reporter assay suggested that the D32Efs*13 23-bp deletion might affect RNA structure. The deleted sequence included the strongest predicted SC35 binding site in an exonic splicing enhancer motif of exon 2 [29, 30] (Supplementary Figure S1). RT-PCR from E10.5 whole embryo RNA with primers in flanking exons identified a major mutant-specific band whose sequence corresponded to splicing of exon 1 to exon 5 in addition to the canonical exon 1 to exon 2 product. Single nucleus RNA sequencing (see below) also identified an exon 1 to exon 5 splice as the more frequent steady-state *Mysm1* RNA in E14.5 cerebellum. The exon1 – exon 5 spliced mRNA encoded a restored open reading frame of 84.6 kD that included each of the annotated MYSM1 protein domains. Together, these data support hypomorphic interpretations of both Y34X (strong) and D32Efs*13 (weak) alleles, with D32Efs*13 having partially restored function through an alternatively spliced mRNA that restored the open reading frame prior to annotated protein domains.

**Supplementary Figure S1.**
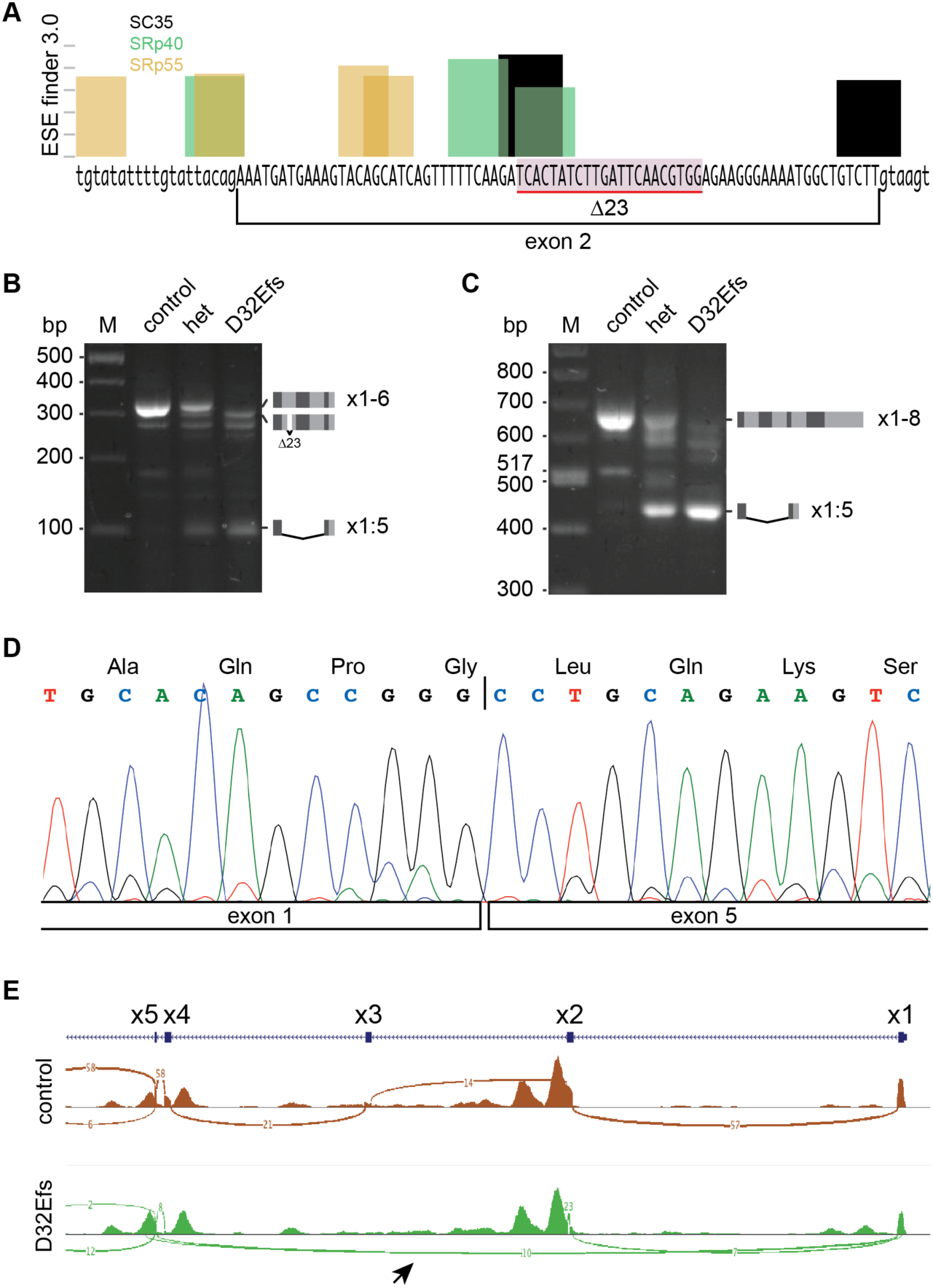
D32Efs*13 mutants express an alternatively spliced RNA that restores reading frame. (**A**) Predicted exonic splicing enhancer protein binding sites in exon 2 showed loss of predicted SC35 and SRp40 sites in the 23-bp deletion. (**B**) RT-PCR with primers in exons 1 and 6 or (**C**) exons 1 and 8 amplified mutant-specific products. (**D**) Sequence of the major mutant-specific RT-PCR product showed precise exon 1 to exon 5 junction. (**E**) Sashimi plot from pseudobulk analysis of single nucleus RNA-seq data also indicated frequent mutant-specific splicing from exon 1 to exon 5 (arrow).

### *Mysm1* mutations disrupt gene expression patterns in multiple cerebellum cell types and decrease proportion of granule cell progenitors

To identify gene regulatory consequences of *Mysm1* mutations in developing cerebellum, we profiled open chromatin and RNA abundance in single nuclei. As morphological consequences are already evident by E16 [31], we focused on E14.5 litters to identify potential antecedent gene regulatory changes in both BKS-*mea^2J^*and FVB–*D32Efs*13* embryos relative to +/+ littermates. Individual cerebellums from four mutant and wild-type littermate pairs, two pairs from each strain, were used for 10X single nucleus multiome analysis. Cluster analysis recovered each of the expected cell types (Figure 4A) from both mutant and control samples (Figure 4B). *Mysm1* was broadly expressed across cell type clusters, indicating that cell autonomy of *mea* phenotypes must be due to different cell type dependencies rather than lineage-restricted expression (Figure 4C). The proportion of nuclei in each cell type differed between genotypes only for granule cell precursors (GCPs, Figure 4D), which comprised a lower percentage of nuclei in mutants for all four littermate pairs (Figure 4E). Among GCPs, UMAP representation was slightly shifted by genotype (Figure 4F), though few individual genes passed statistical threshold at a conservative false discovery rate.

**Figure 4.**
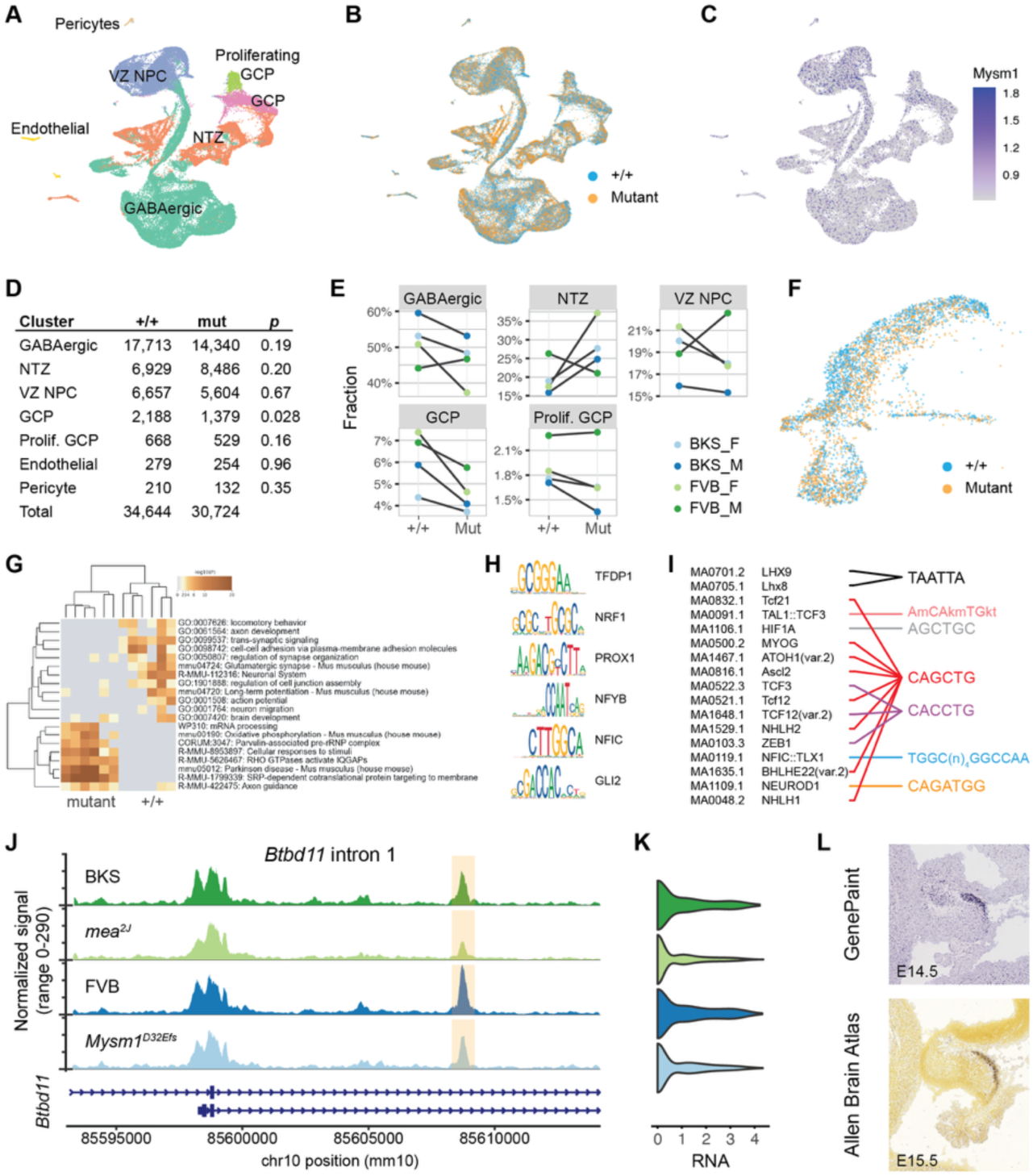
Lower proportion of granule cell precursors, gene expression differences in multiple lineages in E14.5 single-nucleus RNA- and ATAC-seq data. (**A**) UMAP representation of single nuclei clustered by RNA expression reproduced expected cell types. (**B**) Both mutant and control genotypes populated each cluster. (**C**) *Mysm1* was detected above background for cells in all cell-type clusters. (**D**) The proportion of cells in each cluster was significantly different between mutants and controls only for GCP and (**E**) this was consistent for across four littermate pair analyzed from BKS-*mea^2J^*and FVB-*Mysm1^D32Efs^*. (**F**) UMAP distribution of control (blue) and mutant (gold) GCP cells from all eight samples. (**G**) Heat map illustrates RNA expression patterns with gene set differences between *Mysm1* mutant and control nuclei. (**H**) Sequence logos of motifs enriched in control sample ATAC-seq data that have corresponding factors expressed in EGL. (**I**) Motif identifiers and corresponding factors enriched in mutant ATAC-seq data clustered around highly related motifs with shared core sequences across multiple factors expressed in EGL. (**J**) ATAC-seq data showed reduced signal at an intron 1 site in both BKS-*mea^2J^* and FVB-D32Efs cerebellums relative to control littermates. (**K**) Violin plots of snRNA-Seq data show decreased *Btbd11* expression distribution in both mutants. (**L**) *Btbd11* expression localized to the external germinal layer by in situ hybridization in both GenePaint E14.5 and Allen Brain Atlas E15.5 public data [33–35].

To increase power, we considered nuclei for all glutamatergic cells (GCP, proliferating GCP, and nuclear transitory zone) together. For these nuclei, pseudobulk analysis identified 1173 differentially expressed genes at a false discovery rate of 0.05. Functional enrichment analysis showed divergence between genotypes, with control nuclei enriched for gene sets related to cell-cell interactions and synaptic development and mutant nuclei enriched for gene sets related to cell intrinsic functions including mRNA processing, oxidative phosphorylation, and translation (Figure 4G), consistent with the idea that MYSM1 deficiency might delay or diminish maturation in a proportion of cells.

Chromatin accessibility (ATAC-seq) data from the same samples showed 42 chromatin accessibility peaks differed in GCPs by genotype at a false discovery rate of 0.05, 19 with lower signal in mutant than control samples. Of these, 14 differed by genotype in both strain backgrounds and were either physically proximate to or had previously mapped chromatin associations [32] to protein coding genes. To examine potential motif enrichment, we included 943 accessibility peaks that were reduced or enhanced in mutant GCPs relative to littermate controls at a nominal p < 0.01 threshold (540 reduced, 403 enriched). This analysis identified several candidate transcription factor families with one or more members expressed in GCPs of the external germinal layer (EGL) based on public in situ hybridization data [33–35]. Motifs enriched in control samples included sites for factors with an expression bias toward the proliferative outer EGL, including TFDP1 and NFIC (Figure 4H), while motifs enriched in mutant samples included factors associated with both proliferation (ATOH1) and differentiation (LHX9, NEUROD1, NHLH2). Many of these factors had a basic helix-loop-helix (bHLH) domain and clustered into shared core sequences based on their motifs (Figure 4I).

*Btbd11*, which encodes an inhibitory interneuron-specific synaptic protein that promotes glutamatergic signaling [36], provides one example for a GCP-specific gene with significant *Mysm1*-dependence in both chromatin accessibility and expression. *Btbd11* had strong statistical support in both expression and accessibility across all glutamatergic nuclei, with reduced accessibility at a putative enhancer in intron 1 Figure 4J), reduced nuclear RNA expression in mutant samples (Figure 4K) and E14.5 expression predominantly in the external germinal layer (Figure 4L). These expression and chromatin accessibility data together show that *Mysm1* contributes to gene expression patterns across multiple cell types, that dysregulation in the granule cell lineage begins by at least E14.5, that MYSM1 loss can contribute to graded responses across several transcriptional networks, and identifies *Btbd11* as a candidate effector.

### Phylogenetic analysis shows recurrent *Mysm1* loss among Opisthokont lineages

While investigating sequence constraint at the R659-homologous position, we identified apparently recurrent loss of *Mysm1* orthologs among several well-annotated lineages (Figure 5A). We found clear orthologs (best reciprocal BLAST matches to human and mouse MYSM1 and comprising the same Myb-like SANT, SWIRM, and MPN/JAMM domain architecture; Figure 5B) among most animal lineages, including Annelids, Arthropods, Brachiopods, Cnidarians, Echinoderms, Mollusks, as well as four fungal phyla and a few Amoebozoan species. By contrast, *Mysm1* homologs were not found in the most widely studied invertebrate and fungal genomes–Drosophila, Caenorhabditis, and Saccharomyces. While *Mysm1* homologs are present in several insect orders (e.g., Blattodea, Coleoptera, Ephemeroptera) we found no complete homologs among Diptera, Lepidoptera, or Hymenoptera. In contrast to the highly conserved JAMM motif sequences of vertebrate and non-Arthropod invertebrate orthologs, the closest insect homologs had a substitution at the R659-equivalent position in the zinc-coordinating JAMM motif (Figure 5C), potentially suggesting an altered substrate range for the proteolytic domain. On a separate and larger branch, we found no *Mysm1* homologs among any sequenced Nematode species. Despite presence in other fungal phyla, we found no clear homologs among Ascomycota or Basiodiomycota. As these prominent gaps include intensively studied model organisms and several of their wild relatives with multiple, well-annotated genomes, we conclude that the ancestral *Mysm1* gene has been independently lost multiple times in the evolution of both animals and fungi. This pattern of recurrent loss suggests that cellular functions and dependencies can be compensated by activities encoded in other gene genealogies.

**Figure 5.**
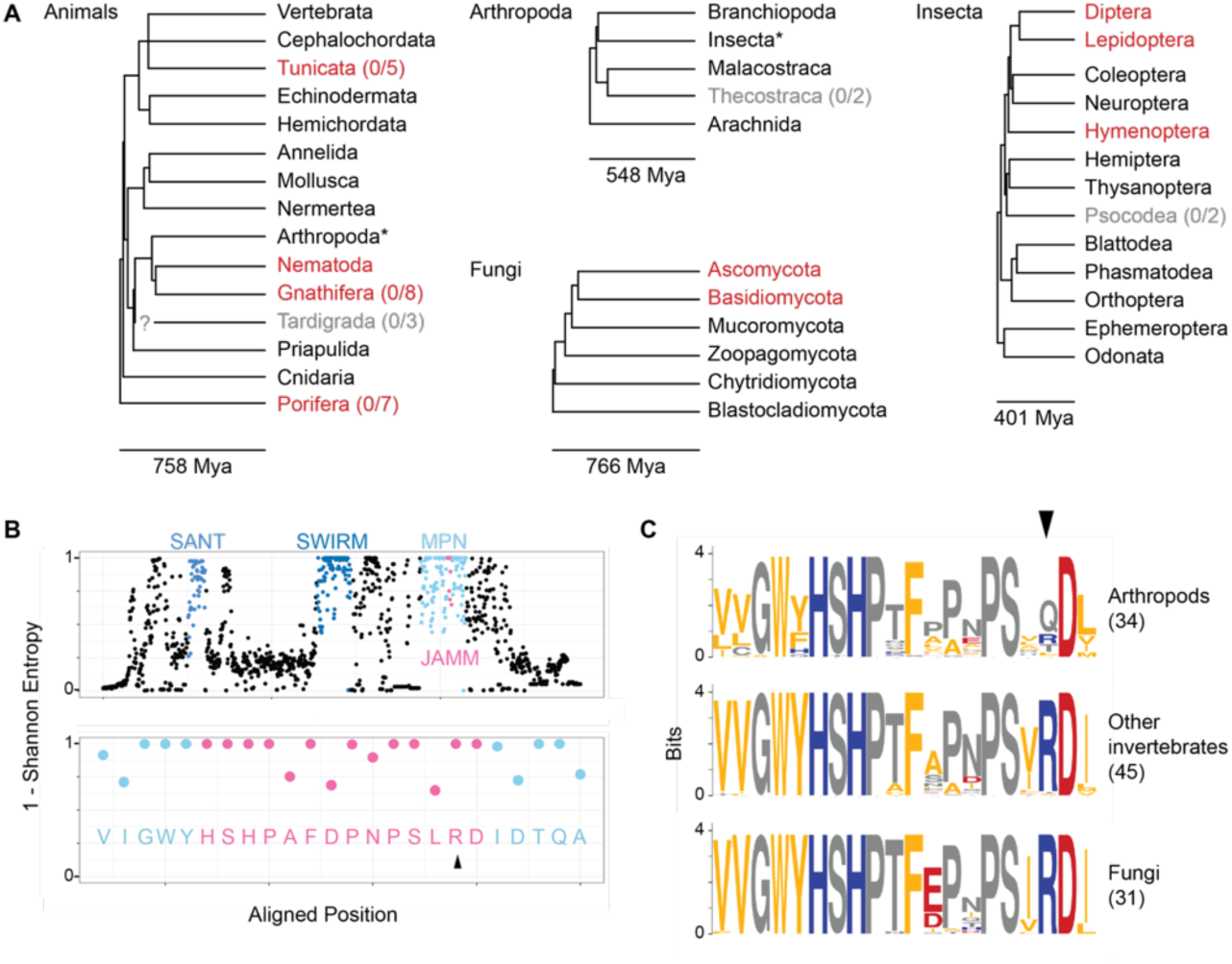
Recurrent Loss of MYSM1 orthologs. (**A**) Phylogenetic trees for animal and fungal phyla, Arthropod classes, and Insect orders using estimated divergence times from TimeTree of Life 5 [37]. Groups with MYSM1 orthologs based on domain architecture [38] and best reciprocal sequence similarity [39] to human and mouse MYSM1 are shown in black. Groups that lack a clear MYSM1 ortholog but include at least 5 species with ANKFN1 [40] or both NXF1 and TULP3 orthologs (Tunicata) are in red. Groups with less evidence for absence in gray. Numbers in parentheses indicate no MYSM1 ortholog given a minimum number of unique species with orthologs for ANKFN1 or TULP3 where the number is less than 10. Asterisks mark groups expanded in an adjacent tree. The position of Tardigrada relative to Arthropoda and Nematoda/Gnathifera is uncertain in TimeTree estimates. (**B**) 1 – Shannon entropy as a measure of site invariance for an alignment of 499 vertebrate (169 mammalian) MYSM1 orthologs, each from a different genus. Canonical domains are highlighted along with the JAMM motif within MPN. Lower panel focused on the JAMM motif within the MPN domain shows that the arginine residue mutated in *mea2J* (arrowhead, equivalent to mouse R659) is invariant among 499 vertebrate orthologs. (**C**) The equivalent position is highly constrained among non-Arthropod invertebrate sequences. Logos show the JAMM motif. While the R659-equivalent position (arrowhead) is nearly invariant among non-Arthropod groups, it is more variable among Arthropods (R only in Daphnia and Arachnids). Y-axis, 0 to 4 bits.

## Discussion

Our data showed that classical *meander tail* (*mea*) mutations are alleles of *Mysm1* and unite a developmental literature for those mutations with more recent literature on MYSM1 cellular and molecular functions. The identification of *mea* with *Mysm1* includes three independent levels of validation: the only plausible mutation in the genetic interval for *mea^J^* is *Mysm1^Y34X^*; a second allele, *mea^2J^*, has a distinct mutation in the same gene, *Mysm1^R659G^*; and new endonuclease-mediated alleles of *Mysm1* failed to complement *mea*. This solves a 50-year mystery on the molecular identity of *meander tail* and expands the range of two previously disparate literatures in confirming broader set of phenotypes in *mea* alleles and adding substantial developmental neurobiology to the MYSM1 literature. The molecular ordering of allele strengths combined with phenotype measures also supports an interpretation that neurological phenotypes are more sensitive to MYSM1 decrement than even the vertebral ankylosing spondylitis characteristic of *meander tail* mutations.

Our data also clarified mechanisms underlying the anterior cerebellum malformation in *mea* mice. Previous work with both chimeras and stem cell transplantation established that the characteristic anterior malformation in *mea* was cell autonomous to granule precursor cells [5, 6]. Identification with *Mysm1* followed by single nucleus sequencing here showed that this reflects a unique requirement for MYSM1 in granule cell precursors rather than restricted expression of the causal gene. While *Mysm1* mutations altered gene expression programs in all cell types we examined, granule cell precursors were uniquely sensitive, with reduced proportion among cell population in each of four littermate pairs at E14.5, two days earlier than previous reports of morphological differences. Thus while our analysis identified a relatively modest number of differential transcripts in GCPs, these are likely to be among the first and most direct gene expression differences in the cell lineage responsible for the malformation. Gene set enrichment showed reduced progression toward neuronal maturation in mutant glutamatergic cells. ATAC-seq data further showed that open chromatin sites preferentially enriched in mutants or controls differed in their enriched transcription factor binding sequence motifs for factors expressed in GCPs. This analysis suggested that motifs in sites with less open chromatin in *Mysm1* mutants may be enriched for factors expressed in less-mature GCPs of the outer EGL. Stronger magnitude effects in R659G (*mea^2J^*), predicted to eliminate metalloprotease activity, than in the hypomorphic D32Efs mutants supports the difference in allele strength seen among gross-level phenotypes. Our single nucleus genomics data showed that MYSM1 affects gene expression across cerebellar cell types, that cell autonomous *mea* phenotypes therefore reflected specific dependency rather than specific expression, showed differential motif enrichment for chromatin accessibility peaks enriched in mutant or control littermate GCPs, and identified candidates for genes affected early in GCP development.

Surprisingly, absence of *Mysm1* orthologues from key model organisms including Drosophila, Caenorhabditis, and Saccharomyces, reflects recurrent loss of *Mysm1* at major lineage branches in both animals and fungi rather than lineage specificity. This recurrence suggests that other deubiquitinases or perhaps some bypass pathway can substitute for MYSM1 activity even in complex animals. These results also reinforce the idea that optimal biomedical progress requires broad consideration of diverse organisms to identify efficient models of specific processes and opportunities for intervention.

## Acknowledgments

We thank the Max Rempel and Trevor Biddle in the UC San Diego Stem Cell Genomics Core at the Sanford Stem Cell Institute for providing PacBio Revio DNA sequencing services. We thank Sang Lee in the Moores Cancer Center Transgenic and CRISPR Mouse Core, supported by NIH grants P30 CA023100 and P30 DK063491, for microinjection of mouse zygotes. We thank Qiongyu Chen in the UCSD Murine Hematology and Coagulation Laboratory for assistance with complete blood counts. We thank the UC San Diego Center for Epigenomics for processing samples for single nucleus assays. This work was supported by NIH grant R21 NS132300 (to BAH) and R35 GM149520 (to CB).

## Methods

### Mice

BKS.Cg–*Dock7^m^* + / + *Lepr^db^* (Stock #000700), BKS.Cg–*mea^J^ Lepr^db^* / + + (Stock #001192), and BKS.Cg–*mea^2J^ Dock7^m^* / + + (Stock #001049) were obtained from live colonies and cryopreserved stocks at The Jackson Laboratories, as were mapping partner strains CAST/EiJ (Stock #000928), A/J (Stock #000646), and FVB/NJ (Stock #001800). Mapping crosses of BKS.Cg–*mea^2J^* x CAST/Ei were performed at the Rockefeller University and National Cancer Institute. Crosses BKS.Cg–*mea^J^* x A/J, BALB/c, and FVB/NJ were performed at UC San Diego. All animal experiments were approved by the Institutional Animal Care and Use Committee of the institution where they were carried out.

#### Phenotype measures

Fresh whole blood was collected into EDTA tubes (BD 365974) after submandibular puncture with a 4 mm lancet. Complete blood counts were obtained with duplicate runs per sample on a HemaVet 950FS (Drew Scientific) programmed with mouse hematology settings. Mutant and control samples were collected and run together but distinct alleles or allelic combinations were run on different days and should not be compared directly. Brain images were obtained from perfusion-fixed samples after bisection at sagittal midline in a brain matrix (Zinc Instruments) as previously described [21, 41]. Mutant and control littermate pairs were processed together. Area measures were made from digital images obtained through a stereoscope after size correction to a ruler in the same image to facilitate comparisons across sample pairs. Folia I-VIa were taken to define the anterior compartment [2].

#### Whole genome sequencing

Spleens from individual animals were frozen in dry ice/ethanol before processing with at the Stem Cell Genomics Core at the Sanford Stem Cell Institute at UC San Diego. Libraries were sequenced to ∼15X coverage with high fidelity (“HiFi”) reads on a PacBio Revio instrument. Barcoded libraries from mutant and control strains were processed together on a single SMRT cell. BAM file alignments were reviewed in the IGV web interface for the critical region of chromosome 4 (89.1 to 102 Mb of mm39 mouse reference).

#### Mysm1 exon sequencing

Primers flanking exons or pairs of exons were designed in Primer3 [42] and used for PCR amplification and Sanger sequencing (Supplementary Table S1). Chromatograms were manually inspected and base calls aligned by BLAT [43] to the mouse reference sequence [24] to identify potential variants. Variant effect predictions (Supplementary Table S2) were conducted using wANNOVAR [26].

#### Genome editing

Pre-assembled high-fidelity SpCas9 ribonucleoprotein complexes were injected into one-cell embryos with minor modifications from our previous studies [41], using the guide sequence ATCACTATCTTGATTCAACG. AltR modified crRNAs, tracrRNA, HiFi SpCas9, Ultramer oligonucleotides as repair templates, HDR Enhancer, and buffers were purchased from IDT (Supplementary Table S3). RNAs were mixed, denatured and annealed prior incubation with diluted Cas9 protein. Final injection mixes were 1.5 µM annealed RNA, 30 ng/µl Cas9, 3.2 µM oligonucleotides and 1 µM (exon 6) or 10µM (exon 2) HDR Enhancer v.2. Targeted exons were amplified by PCR from tail DNA of resulting G0 pups and sequenced as above.

#### Western blots

Dissected tissues were disrupted in RIPA buffer (Teknova, R3792) with protease inhibitors (Sigma-Aldrich P8340), sonicated, and concentration measured with a bicinchoninic acid (BCA) assay (Pierce, 23225). 20 µg of protein from mutant and control littermate samples were fractionated on 8% acrylamide gels and electrophoretically transferred to nitrocellulose membranes (Bio-Rad, 1620115). After blocking with 5% non-fat dry milk in PBST, membranes were exposed to rabbit anti-MYSM1 antibody (Invitrogen PA5-102696, raised against residues 645-695 relative to Q5VVJ2, including the JAMM motif) and mouse anti-GAPDH (GeneTex GTX627408) in blocking solution for 1 hour at room temperature, washed 4x in blocking solution, then exposed to goat anti-rabbit IR-680 (Li-Cor 92568071) and goat anti-mouse IR-800 (Li-Cor 92632210) dye-conjugated secondary antibodies for 1 hour at room temperature, and washed once in blocking solution, twice in PBST and once in PBS. Fluorescence was quantified on a LI-COR Odyssey instrument with ImageStudio software (LiCOR).

#### Reporter assays

Mouse and human *Mysm1* fragments were fused to a Nano-luciferase reporter in-frame with respect to wild-type MYSM1 in a dual reporter plasmid we recently developed [21]. Briefly, the *Zfp423* 5’ fragment from pZNpF-wt (Addgene #245364) was excised by digestion with EcoRI-HF and BamHI-HF (New England Biolabs) and replaced by chemically synthesized mouse *Mysm1* or human *MYSM1* fragments (Twist Bioscience) using a HiFi DNA Assembly Cloning Kit (New England Biolabs). All plasmids were fully sequenced (Plasmidsaurus or Quintara Biosciences) before use. P19 cells were transfected in 96-well plates with at least four independent DNA preparations per construct and harvested the following day. Reporter expression was quantifies using a Nano-Glo dual-luciferase reporter assay system (Promega) according to the manufacturer’s protocol as previously adapted [21] to a Tecan Spark plate reader. In luminescence mode (360-700 nm wavelength, 500 ms). Luminescence measures were adjusted to negative control plasmids for each reporter. Relative Response Ratio (RRR) was calculated from the ratio of adjusted nano to firefly luciferase values; RRR = (sample ratio—negative control ratio)/(positive control ratio—negative control ratio). Log_2_-transformed values of RRR were used for visualization and ANOVA models.

#### RT-PCR

E10.5 whole-embryo RNA was extracted with Trizol reagent, reverse transcribed with SuperScript II (Invitrogen), and subjected to 35 cycles of amplification with primers pairs in exons 1 and 6 or exons 1 and 8 (Supplementary Table S1). Gel-isolated bands were used as templates for Sanger sequencing from the reverse primer to allow accurate coverage of the exon 1 splice junctions.

#### Single nucleus genomic assays

Timed matings were set up and noon of the following day was designated as embryonic day (E)0.5. Cerebellum from E14.5 litters were manually dissected in ice-cold PBS and flash-frozen in either a dry ice methanol bath (1 pair) or liquid nitrogen (3 pairs). Tails were used for PCR-based genotyping with allele-specific assays. Nuclei were prepared and processed through 10X Multiome ATAC + Gene Expression in the UC San Diego Center for Epigenomics. Cell type expression inferred from clustering was confirmed for several examples from public in situ hybridization databases [33–35]. *Btbd11* examples shown were accessed from GenePaint (https://gp3.mpg.de/results/btbd11) and Allen Developing Mouse Brain (https://developingmouse.brain-map.org/experiment/show/10005457).

#### Single nucleus data analysis

Raw sequencing reads were processed using Cell Ranger ARC 2.0.2 with the mm10 2020-A transcriptome reference from 10X Genomics. Following sample-level counting, all outputs were aggregated with library normalization mode “none.” Cells with greater than 10% ribosomal or mitochondrial RNA reads were filtered out, as were cells with fewer than 500 RNA reads. Possible doublets were removed using scDblFinder [44] on each sample independently, with random artificial doublet generation and default parameters. Within scDblFinder, the AMULET [45] implementation was also used to identify likely doublets based on greater than expected ATAC-seq coverage. Within Seurat [46], gene expression processing was carried out using default parameters for the NormalizeData(), FindVariableFeatures(), ScaleData(), RunPCA(), and FindNeighbors() functions. Louvain clustering with a resolution of 0.5 yielded 19 total clusters. UMAP generation was performed using the first 50 principal components. To calculate differential abundance of cell types across genotypes, the propeller method [47] was used to transform cell type proportions. The significance of these values was then assessed using a paired t-test that matched mice according to sex and strain background. A summary of functional enrichment across cell types was carried out using differentially expressed genes between genotypes based on a pseudobulk analysis of the six largest Louvain clusters. Differential chromatin accessibility and motif enrichment were calculated using Signac [48]. GCP-enriched peaks were identified using the logistic regression method in FindMarkers(), with cell-level ATAC-seq counts specified as a latent variable. The 16,866 peaks with adjusted p-value less than 0.05 were then used for motif finding against the default 40,000 GC-matched background features. Differentially accessible peaks between genotypes within the GCP population were identified using the same approach. Motif enrichment for each genotype was calculated using peaks passing a less stringent 0.01 raw p-value threshold. The single-nucleus RNA-and ATAC-seq data have been deposited in NCBI GEO with the accession number GSE318479.

#### Conservation analyses

1709 pre-computed vertebrate orthologs were obtained using the NCBI Datasets command line tool. A single representative sequence was manually selected from each genus and any remaining identical protein sequences were removed. Three manually-curated orthologs (*Lampetra*, *Umbra*, and *Lota*) were added from BLAST searches of early-branching and under-sampled orders to produce a curated set of 499 unique vertebrate MYSM1 sequences (Supplementary Table S4). Multiple sequence alignment was generated using CLUSTAL-Omega [49] from its web interface at the European Bioinformatics Institute [50] under default parameters. Each position in the alignment was scored for Shannon entropy as a measure of residue homogeneity [51] from a previously described web tool (https://compbio.cs.princeton.edu/conservation/score.html) [52] and resulting scores were plotted using ggplot2 [53]. Invertebrate (Supplementary Table S5) and fungal (Supplementary Table S6) homologs were manually curated from iterative and reciprocal BLAST searches to identify homologs where automated annotation terms were less accurate due to reduced identity and matches to non-MYSM1 proteins with overlapping domain architecture but orthologous to other mammalian proteins. Reciprocal best matches to vertebrate MYSM1 proteins with conserved domain architecture were taken as presumptive orthologs. Absence of an MYSM1 ortholog was scored for species with well-annotated genomes that included orthologs for NXF1 and either ANKFN1, which is poorly annotated in incomplete genomes [40], or TUBBY-like proteins as controls for annotation. Lineages were deemed likely to have lost MYSM1 if they included such genomes from at least 5 distinct species and no evidence for MYSM1.

## Cited Literature

1. Hollander, W.F. and K.S. Waggie, Meander tail: a recessive mutant located in chromosome 4 of the mouse. J Hered, 1977. 68(6): p. 403–6.

2. Ross, M.E., et al., Meander tail reveals a discrete developmental unit in the mouse cerebellum. Proc Natl Acad Sci U S A, 1990. 87(11): p. 4189–92.

3. Napieralski, J.A. and L.M. Eisenman, Further evidence for a unique developmental compartment in the cerebellum of the meander tail mutant mouse as revealed by the quantitative analysis of Purkinje cells. J Comp Neurol, 1996. 364(4): p. 718–28.

4. Hamre, K.M. and D. Goldowitz, Analysis of gene action in the meander tail mutant mouse: examination of cerebellar phenotype and mitotic activity of granule cell neuroblasts. J Comp Neurol, 1996. 368(2): p. 304–15.

5. Hamre, K.M. and D. Goldowitz, meander tail acts intrinsic to granule cell precursors to disrupt cerebellar development: analysis of meander tail chimeric mice. Development, 1997. 124(21): p. 4201–12.

6. Rosario, C.M., et al., Differentiation of engrafted multipotent neural progenitors towards replacement of missing granule neurons in meander tail cerebellum may help determine the locus of mutant gene action. Development, 1997. 124(21): p. 4213–24.

7. Zhu, P., et al., A histone H2A deubiquitinase complex coordinating histone acetylation and H1 dissociation in transcriptional regulation. Mol Cell, 2007. 27(4): p. 609–21.

8. Zhou, W., et al., Histone H2A monoubiquitination represses transcription by inhibiting RNA polymerase II transcriptional elongation. Mol Cell, 2008. 29(1): p. 69–80.

9. Jiang, X.X., et al., Control of B cell development by the histone H2A deubiquitinase MYSM1. Immunity, 2011. 35(6): p. 883–96.

10. Nijnik, A., et al., The critical role of histone H2A-deubiquitinase Mysm1 in hematopoiesis and lymphocyte differentiation. Blood, 2012. 119(6): p. 1370–9.

11. Liakath-Ali, K., et al., Novel skin phenotypes revealed by a genome-wide mouse reverse genetic screen. Nat Commun, 2014. 5: p. 3540.

12. Panda, S., J.A. Nilsson, and N.O. Gekara, Deubiquitinase MYSM1 Regulates Innate Immunity through Inactivation of TRAF3 and TRAF6 Complexes. Immunity, 2015. 43(4): p. 647–59.

13. Alsultan, A., et al., MYSM1 is mutated in a family with transient transfusion-dependent anemia, mild thrombocytopenia, and low NK- and B-cell counts. Blood, 2013. 122(23): p. 3844–5.

14. Le Guen, T., et al., An in vivo genetic reversion highlights the crucial role of Myb-Like, SWIRM, and MPN domains 1 (MYSM1) in human hematopoiesis and lymphocyte differentiation. J Allergy Clin Immunol, 2015. 136(6): p. 1619–1626 e5.

15. Bahrami, E., et al., Myb-like, SWIRM, and MPN domains 1 (MYSM1) deficiency: Genotoxic stress-associated bone marrow failure and developmental aberrations. J Allergy Clin Immunol, 2017. 140(4): p. 1112–1119.

16. Sweet, H.O. and M.T. Davisson, Remutations at The Jackson Laboratory. Mouse Genome, 1995. 93(4): p. 1030–1034.

17. Sweet, H.O., Remutations at The Jackson Laboratory. Mouse Genome, 1993. 91(4): p. 862–865.

18. Fletcher, C., D.J. Norman, and N. Heintz, Genetic mapping of meander tail, a mouse mutation affecting cerebellar development. Genomics, 1991. 9(4): p. 647–55.

19. Baudat, F., et al., PRDM9 is a major determinant of meiotic recombination hotspots in humans and mice. Science, 2010. 327(5967): p. 836–40.

20. Parvanov, E.D., P.M. Petkov, and K. Paigen, Prdm9 controls activation of mammalian recombination hotspots. Science, 2010. 327(5967): p. 835.

21. Concepcion, D., et al., Nonequivalence of Zfp423 premature termination codons in mice. Genetics, 2025. 231(2).

22. Kozak, M., Constraints on reinitiation of translation in mammals. Nucleic Acids Res, 2001. 29(24): p. 5226–32.

23. Sherlock, M.E., et al., Principles, mechanisms, and biological implications of translation termination-reinitiation. RNA, 2023. 29(7): p. 865–884.

24. Raney, B.J., et al., The UCSC Genome Browser database: 2024 update. Nucleic Acids Res, 2024. 52(D1): p. D1082–D1088.

25. Karczewski, K.J., et al., Variation across 141,456 human exomes and genomes reveals the spectrum of loss-of-function intolerance across human protein-coding genes. bioRxiv, 2019: p. 531210.

26. Yang, H. and K. Wang, Genomic variant annotation and prioritization with ANNOVAR and wANNOVAR. Nat Protoc, 2015. 10(10): p. 1556–66.

27. Cook, A.G., et al., Cell division angle predicts the level of tissue mechanics that tune the amount of cerebellar folding. Development, 2024. 151(3).

28. Concepcion, D., et al., Nonequivalence of Zfp423 premature termination codons in mice. Genetics, 2025: p. iyaf164.

29. Cartegni, L., et al., ESEfinder: A web resource to identify exonic splicing enhancers. Nucleic Acids Res, 2003. 31(13): p. 3568–71.

30. Smith, P.J., et al., An increased specificity score matrix for the prediction of SF2/ASF-specific exonic splicing enhancers. Hum Mol Genet, 2006. 15(16): p. 2490–508.

31. Napieralski, J.A. and L.M. Eisenman, Developmental analysis of the external granular layer in the meander tail mutant mouse: do cerebellar microneurons have independent progenitors? Dev Dyn, 1993. 197(4): p. 244–54.

32. Gorkin, D.U., et al., An atlas of dynamic chromatin landscapes in mouse fetal development. Nature, 2020. 583(7818): p. 744–751.

33. Visel, A., C. Thaller, and G. Eichele, GenePaint.org: an atlas of gene expression patterns in the mouse embryo. Nucleic Acids Res, 2004. 32(Database issue): p. D552–6.

34. Diez-Roux, G., et al., A high-resolution anatomical atlas of the transcriptome in the mouse embryo. PLoS Biol, 2011. 9(1): p. e1000582.

35. Henry, A.M. and J.G. Hohmann, High-resolution gene expression atlases for adult and developing mouse brain and spinal cord. Mamm Genome, 2012. 23(9-10): p. 539–49.

36. Bygrave, A.M., et al., Btbd11 supports cell-type-specific synaptic function. Cell Rep, 2023. 42(6): p. 112591.

37. Kumar, S., et al., TimeTree 5: An Expanded Resource for Species Divergence Times. Mol Biol Evol, 2022. 39(8).

38. Letunic, I., S. Khedkar, and P. Bork, SMART: recent updates, new developments and status in 2020. Nucleic Acids Res, 2021. 49(D1): p. D458–D460.

39. Altschul, S.F., et al., Basic local alignment search tool. J Mol Biol, 1990. 215(3): p. 403–10.

40. Zhang, S., et al., Nmf9 Encodes a Highly Conserved Protein Important to Neurological Function in Mice and Flies. PLoS Genet, 2015. 11(7): p. e1005344.

41. Deshpande, O., et al., ZNF423 patient variants, truncations, and in-frame deletions in mice define an allele-dependent range of midline brain abnormalities. PLoS Genet, 2020. 16(9): p. e1009017.

42. Rozen, S. and H. Skaletsky, Primer3 on the WWW for general users and for biologist programmers. Methods Mol Biol, 2000. 132: p. 365–86.

43. Kent, W.J., BLAT--the BLAST-like alignment tool. Genome Res, 2002. 12(4): p. 656–64.

44. Germain, P.L., et al., Doublet identification in single-cell sequencing data using scDblFinder. F1000Res, 2021. 10: p. 979.

45. Thibodeau, A., et al., AMULET: a novel read count-based method for effective multiplet detection from single nucleus ATAC-seq data. Genome Biol, 2021. 22(1): p. 252.

46. Butler, A., et al., Integrating single-cell transcriptomic data across different conditions, technologies, and species. Nat Biotechnol, 2018. 36(5): p. 411–420.

47. Phipson, B., et al., propeller: testing for differences in cell type proportions in single cell data. Bioinformatics, 2022. 38(20): p. 4720–4726.

48. Stuart, T., et al., Single-cell chromatin state analysis with Signac. Nat Methods, 2021. 18(11): p. 1333–1341.

49. Sievers, F., et al., Fast, scalable generation of high-quality protein multiple sequence alignments using Clustal Omega. Mol Syst Biol, 2011. 7: p. 539.

50. Madeira, F., et al., The EMBL-EBI Job Dispatcher sequence analysis tools framework in 2024. Nucleic Acids Res, 2024. 52(W1): p. W521–W525.

51. Valdar, W.S., Scoring residue conservation. Proteins, 2002. 48(2): p. 227–41.

52. Capra, J.A. and M. Singh, Predicting functionally important residues from sequence conservation. Bioinformatics, 2007. 23(15): p. 1875–82.

53. Wickham, H., ggplot2: Elegant Graphics for Data Analysis. 2016: Springer-Verlag New York.

